# Broad antifungal resistance mediated by RNAi-dependent epimutation in the basal human fungal pathogen *Mucor circinelloides*

**DOI:** 10.1101/526459

**Authors:** Zanetta Chang, R. Blake Billmyre, Soo Chan Lee, Joseph Heitman

**Affiliations:** Department of Molecular Genetics and Microbiology, Duke University, Duke University Medical Center, Durham, North Carolina, United States of America; Stowers Institute for Medical Research, Kansas City, Missouri, United States of America; South Texas Center for Emerging Infectious Diseases (STCEID), Department of Biology, University of Texas, San Antonio, San Antonio, Texas, United States of America

## Abstract

Mucormycosis - an emergent, deadly fungal infection - is difficult to treat, in part because the causative species demonstrate broad clinical antifungal resistance. However, the mechanisms underlying drug resistance in these infections remain poorly understood. Our previous work demonstrated that one major agent of mucormycosis, Mucor circinelloides, can develop resistance to the antifungal agents FK506 and rapamycin through a novel, transient RNA interference-dependent mechanism known as epimutation. Epimutations silence the drug target gene and are selected by drug exposure; the target gene is re-expressed and sensitivity is restored following passage without drug. This silencing process involves generation of small RNA (sRNA) against the target gene via core RNAi pathway proteins. To further elucidate the role of epimutation in the broad antifungal resistance of Mucor, epimutants were isolated that confer resistance to another antifungal agent, 5-fluoroorotic acid (5-FOA). We identified epimutant strains that exhibit resistance to 5-FOA without mutations in PyrF or PyrG, enzymes which convert 5-FOA into the active toxic form. Using sRNA hybridization as well as sRNA library analysis, we demonstrate that these epimutants harbor sRNA against either *pyrF* or *pyrG*, and further show that this sRNA is lost after reversion to drug sensitivity. We conclude that epimutation is a mechanism capable of targeting multiple genes, enabling Mucor to develop resistance to a variety of antifungal agents. Elucidation of the role of RNAi in epimutation affords a fuller understanding of mucormycosis. Furthermore, it improves our understanding of fungal pathogenesis and adaptation to stresses, including the evolution of drug resistance.

**Author Summary:** The emerging infection mucormycosis causes high mortality in part because the major causative fungi, including *Mucor circinelloides*, are resistant to most clinically available antifungal drugs. We previously discovered an RNA interference-based resistance mechanism, epimutation, through which *M*. *circinelloides* develops transient resistance to the antifungal agent FK506 by altering endogenous RNA expression. We further characterize this novel mechanism by isolating epimutations in two genes that confer resistance to another antifungal agent, 5-fluoroorotic acid. Thus, we demonstrate epimutation can induce resistance to multiple antifungals by targeting a variety of genes. These results reveal epimutation plays a broad role enabling rapid and reversible fungal responses to environmental stresses, including drug exposure, and controlling antifungal drug resistance and RNA expression. As resistance to antifungals emerges, a deeper understanding of the causative mechanisms is crucial for improving treatment.

## Introduction

Mucormycosis, an emerging fungal infection, is notable for very high mortality, ranging from 50% for rhino-orbital-cerebral infections to 90% in disseminated infections [1]. Mucormycosis primarily affects immunocompromised patients: most commonly patients with diabetes, followed by those with hematologic cancers, prior organ transplants, trauma, and iron overload disorders [2, 3]. The increasing prevalence of these immunosuppressive disorders may explain the rising incidence of mucormycosis. Another major problem is that treatment options are very limited, with first-line therapy consisting of surgical debridement combined with amphotericin B or isavuconazole, the only FDA-approved antifungal agents for mucormycosis [4-6]. Even after recovery patients often suffer from permanent disfigurement.

The etiologic causes of mucormycosis are the Mucoralean fungi, of which the three most common infectious genera are *Rhizopus, Mucor*, and *Lichtheimia* [7]. Of these genera, *Mucor* has served as a model organism in various aspects of fungal biology (e.g. RNAi biology, virulence, and light sensing), and the scientific community has developed a set of tools for genetic manipulation [8-12]. Despite this knowledge base, many gaps remain in our understanding of the pathogenesis of *Mucor* as well as the biology of all Mucoralean species. For example, the broad, intrinsic antifungal resistance common to Mucoralean fungi results in limited treatment options and may contribute to the high mortality associated with mucormycosis, yet the mechanisms underlying this resistance remain largely uncharacterized. Our laboratory previously identified a form of drug resistance in *Mucor* that is dependent upon endogenous RNA interference (RNAi), referred to as epimutation [13, 14].

RNAi is a mechanism that targets specific mRNA transcripts and inactivates them through either mRNA degradation or inhibition of translation. The first description of RNAi in fungi was quelling, a mechanism for silencing repetitive sequences and transposons in the model fungus *Neurospora crassa* [15]. Later, RNAi was fully characterized in the nematode *Caenorhabditis elegans* [16], and has since been shown to be conserved throughout many eukaryotic lineages including a variety of fungi [17]. Many other forms of RNAi have since been characterized in fungi, including meiotic silencing by unpaired DNA in *Neurospora*, sex-induced silencing in *Cryptococcus neoformans*, and heterochromatin formation in *Schizosaccharomyces pombe* [18-21]. RNA-based control of fungal drug sensitivity was previously described in *S*. *pombe*, where a long non-coding RNA has been shown to epigenetically repress transcription of a permease and, therefore, decrease global drug sensitivity [22]. However, no RNAi-mediated form of drug resistance was described prior to our previous finding of epimutation in *Mucor* [13, 14]. In the Mucoralean fungi, RNAi machinery is conserved and functions to trigger silencing in *Mucor circinelloides* and *Rhizopus delemar/Rhizopus oryzae* [8, 23, 24]. Thus, *Mucor* serves as a model fungus for the study of RNAi and epimutation.

The novel mechanism of epimutation involves the intrinsic RNAi silencing pathway, which transiently suppresses expression of fungal drug target genes. Epimutants in *Mucor* were previously identified that confer resistance to the antifungal agents FK506 and rapamycin. These epimutants harbor antisense small RNAs (sRNA) specific to the *fkbA* gene that trigger mRNA degradation and thereby prevent production of the drug target FKBP12 [13]. Epimutation is transient; after passage without FK506 drug selection, mRNA expression recovered gradually until *fkbA* mRNA returned to wild-type expression levels and, conversely, expression of the *fkbA*-specific sRNA was lost. Epimutation in *Mucor* requires multiple canonical RNAi proteins, including Dicer (Dcl1 and Dcl2), Argonaute (Ago1), and RNA-dependent RNA polymerase (RdRP2). Deletion of genes encoding these RNAi components in *M*. *circinelloides* led to an inability to form epimutants, showing that epimutation is dependent upon the RNAi pathway. Interestingly, deletion of two other RNA-dependent RNA polymerases, RdRP1 or RdRP3, or the RNAi pathway component R3B2, led to a significantly higher rate of epimutation, suggesting these components play an inhibitory role [14]. Taken together, these findings reveal the intrinsic RNAi pathway in *Mucor* can suppress drug target expression in a reversible fashion.

We report here the identification of epimutants resistant to an additional antifungal, the laboratory agent 5-fluoroorotic acid (5-FOA). 5-FOA is converted into a toxin by action of orotate phosphoribosyltransferase (PyrF) and orotidine-5'-monophosphate decarboxylase (PyrG), two enzymes in the uracil biosynthetic pathway. Antifungal resistance is evoked by selective generation of sRNA against either *pyrF* and *pyrG*. Similar to previous observations with FK506-resistant epimutants, sRNA generation in 5-FOA-resistant epimutants is transient and lost after passage in the absence of drug selection, or when the epimutants are grown in conditions lacking uracil. These observations build on our prior findings to establish that epimutation is a general phenomenon that can affect multiple genetic loci in *Mucor* and induce resistance to multiple antifungal agents. The transient nature of epimutation allows for rapid adaptation through generation of phenotypic diversity in response to a variety of stresses, such as drug stress or auxotrophy. These findings advance our understanding of the genetic and molecular basis for antimicrobial drug resistance with implications for other pathogenic microbes with active RNAi pathways.

## Results

### Epimutation induces transient resistance to 5-FOA

Our previous work identified epimutation as a novel mechanism of antifungal resistance, but we had only studied sRNAs generated against a single locus, *fkbA* [13]. To determine the broader scope epimutation might play in *Mucor* drug resistance, we generated epimutations against another antifungal compound. The well-characterized laboratory antifungal agent 5-fluoroorotic acid (5-FOA) possesses efficacy against *Mucor* and is used as a tool for genetic manipulation [12, 25-27]. The genes responsible for 5-FOA toxicity in *Mucor* encode orotate phosphoribosyltransferase (*pyrF*) and orotidine-5’-monophosphate decarboxylase (*pyrG*) [28, 29]. PyrF and PyrG are responsible for the conversion of 5-FOA, a prodrug, into 5-fluorouracil, which serves as a toxic nucleotide analog. Therefore, a loss-of-function mutation in either of these two genes confers resistance. Because these genes also play roles in the pyrimidine synthesis pathway, *pyrF* or *pyrG* mutation also causes uracil auxotrophy. The clear understanding of the mechanisms and targets of 5-FOA simplified the process of screening for epimutants.

To increase the possibility of isolating epimutants we performed initial screens in two RNAi-deficient backgrounds, *rdrp1*Δ and *rdrp3*Δ, which demonstrated an enhanced rate of epimutation in previous studies of FK506-resistant epimutation [14]. These *rdrp1*Δ and *rdrp3*Δ mutant strains contain two copies of *pyrG*. The original *pyrG* locus contains a known point mutation, G413A, which confers 5-FOA resistance. Due to the limited selectable markers available for *Mucor*, a functional copy of *pyrG* was subsequently inserted into either the *rdrp1* or *rdrp3* locus to generate the RNAi mutant strains. Therefore, to sequence and identify *pyrG* mutations in RNAi mutant strains, we specifically amplified the copy of *pyrG* inserted in either *rdrp1* or *rdrp3* using the appropriate locus-specific primers (S1 Table). Of note, all of the *pyrG* mutations found in this study match the original mutation seen in the endogenous mutant *pyrG* locus (S2 Table). This is most likely due to gene conversion from the endogenous locus, indicating a higher rate of gene conversion when compared to *de novo* mutation. This phenomenon may have contributed to the relatively low frequency of isolation of *pyrG* epimutants.

To derive 5-FOA-resistant isolates, *rdrp1*Δ and *rdrp3*Δ strains were grown in the presence of 5-FOA in media supplemented with uridine and uracil. Under these conditions the strains, initially sensitive, grew as abnormal, stunted hyphae; but after approximately two weeks of incubation patches of resistant filamentous growth were isolated and analyzed. Two *pyrF* epimutants (designated as strains E1 and E2) were isolated from an *rdrp3* mutant strain. In addition, four *pyrF* epimutants (including strains E3 and E5) and one *pyrG* epimutant (E4) were isolated in an *rdrp1* mutant strain, a second strain with enhanced rates of epimutation [13]. Based on sRNA hybridization analysis, representative epimutants express antisense sRNA against either the *pyrF* or *pyrG* locus, but not both (Fig. 1a, b). No 5-FOA-resistant epimutant strains were identified in wild-type strains R7B (*M*. *circinelloides* f. *lusitanicus*) or 1006PhL (*M*. *circinelloides* f. *circinelloides*), or in an *r3b2Δ* strain, mutated for a different RNAi component (S2 Table). Interestingly, rates of 5-FOA-resistant epimutation in all strains tested were decreased compared to the rates seen in the initial report of FK506-resistant epimutants (Table 1).

**Table 1.** Rates of *pyrF* and *pyrG* epimutation by background strain

|  |  | Total | Epimutant | Mutant | Unknown |
| --- | --- | --- | --- | --- | --- |
| *rdrp3Δ* | Number of isolates | 14 | 2 | 0 | 12 |
|  | Percentage |  | 14.3% | 0% | 85.7% |
| *rdrp1Δ* | Number of isolates | 27 | 5 | 20 | 2 |
|  | Percentage |  | 18.5% | 74.1% | 7.4% |
| *r3b2Δ* | Number of isolates | 14 | 0 | 12 | 2 |
|  | Percentage |  | 0% | 85.7% | 14.3% |
| R7B | Number of isolates | 2 | 0 | 0 | 2 |
|  | Percentage |  | 0% | 0% | 100.0% |
| 1006PhL | Number of isolates | 61 | 0 | 28 | 33 |
|  | Percentage |  | 0% | 45.9% | 54.1% |

**Fig 1.**
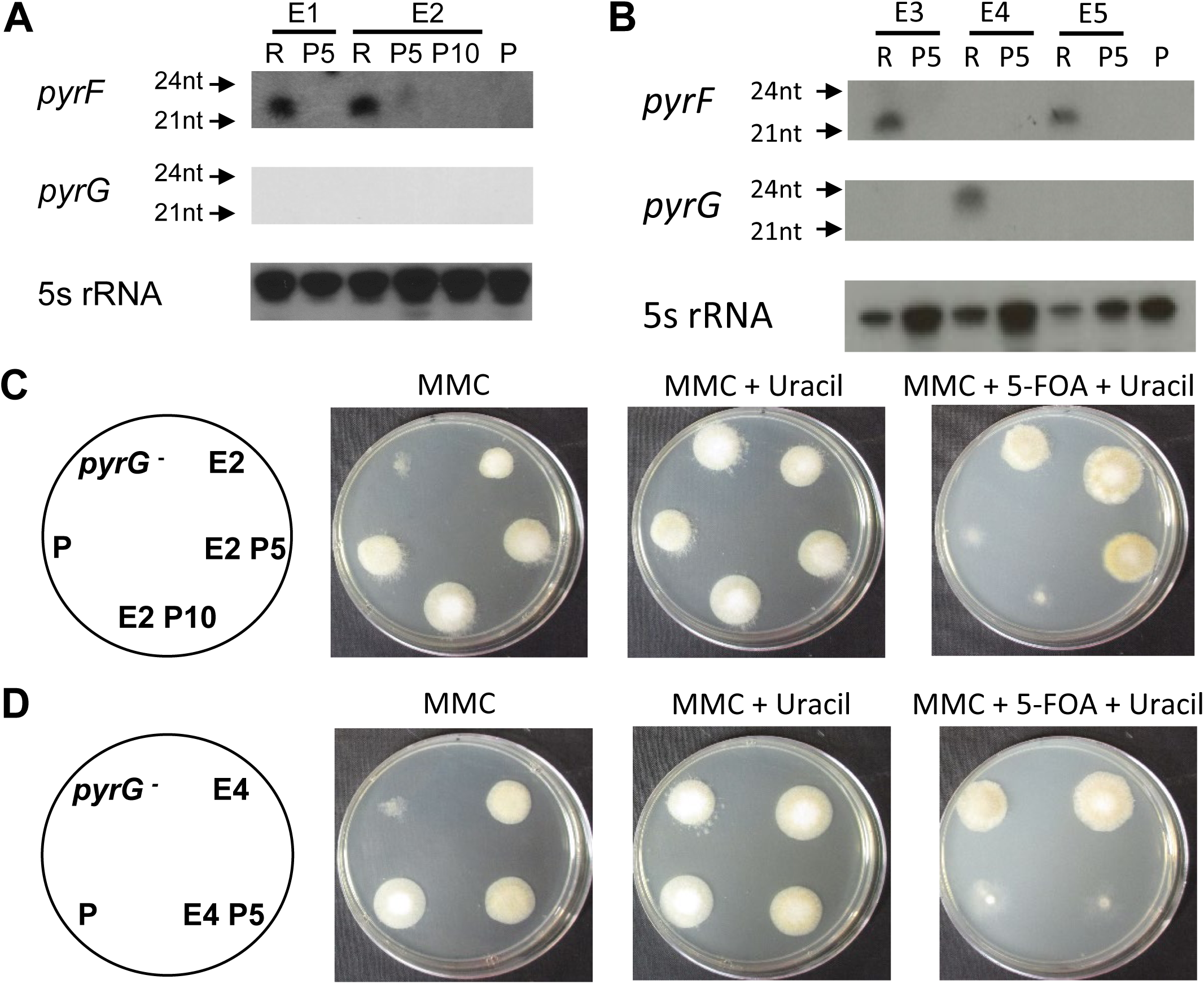
sRNA hybridization and phenotypic analysis of 5-FOA-resistant epimutants. **(A)** sRNA hybridization of epimutants and revertants from an *rdrp3* mutant background, before (R – resistant) and after 5 (P5) or 10 (P10) passages without selection. P, *rdrp3Δ* parental strain (MU439). Blots were hybridized with antisense-specific probes against *pyrF, pyrG*, or 5S rRNA (loading control). **(B)** sRNA blot of epimutants and revertants from an *rdrp1* mutant background, before (R – resistant) and after 5 passages without selection (P5). P, *rdrp1Δ* parental strain (MU419). Blots were hybridized with antisense-specific probes against *pyrF, pyrG* or 5S rRNA (loading control). **(C)** Phenotypic analysis of one representative epimutant, before and after reversion. A *pyrF* epimutant (E2) is shown before and after 5 (P5) and 10 (P10) passages without selection, grown on MMC media, MMC supplemented with uridine and uracil, and MMC supplemented with 5-FOA, uridine, and uracil. *pyrG*^*-*^, a known mutant of *pyrG*, served as a negative control; P, *rdrp3Δ* parental strain (MU439). **(D)** A *pyrG* epimutant (E4) is shown before and after 5 (P5) passages without selection, grown on MMC, MMC supplemented with uracil, and MMC supplemented with 5-FOA and uracil. P, *rdrp1Δ* parental strain (MU419).

All epimutant strains were stably 5-FOA-resistant when maintained under drug selection conditions. However, following passage on media lacking 5-FOA, all five strains reverted to a 5-FOA sensitive phenotype. To determine 5-FOA sensitivity, epimutant and passaged strains were plated on MMC media without uracil, MMC with uracil supplementation, and MMC with 5-FOA and uracil for phenotypic analysis (Fig. 1c, d). Uracil auxotrophic strains with known mutations, such as the *pyrG-* mutant strain, are unable to grow robustly on MMC alone. In contrast, the epimutant strains were able to grow to some extent on MMC. Epimutant E2 shows qualitatively reduced growth on agar plates relative to the parental strain, while epimutant E4 shows growth indistinguishable from the parental strain. This suggests that epimutants placed in auxotrophic conditions may still be able to synthesize uracil at a low level; or, alternatively, that the epimutation has begun to revert toward wild-type when epimutant spores are incubated on MMC. Complete reversion of epimutant strains - loss of 5-FOA resistance and wild-type rates of growth on MMC - was observed for *pyrF* epimutants E1, E3, and E5, as well as *pyrG* epimutant E4 after five passages (Fig. 1d). The *pyrF* epimutant E2 demonstrated only partial reversion to drug sensitivity after five passages but complete reversion after ten (Fig. 1c). sRNA was isolated from these reverted strains after five or ten passages and sRNA hybridization was performed. Strains E1, E3, E4, and E5 demonstrated a complete loss of *pyrF* or *pyrG* sRNA after five passages, corresponding with their phenotypic reversion; likewise, strain E2 demonstrated a reduction of *pyrF* sRNA after five passages and complete loss after ten passages (Fig. 1a, b).

### 5-FOA-resistant epimutants express sRNA specific to the *pyrF* or *pyrG* locus

sRNA libraries were generated from *pyrF* epimutants (E1, E2) and the *pyrG* epimutant (E4) as well as their corresponding revertants, and these libraries were sequenced via Illumina. Epimutation induced a significant increase in both sense and antisense sRNAs against *pyrF* or *pyrG* in their respective epimutants. For *pyrF*, which contains no introns, these sRNAs were distributed across the ORF (Fig. 2a). sRNAs expressed against *pyrG* were localized specifically to the exons (Fig. 2b). In both cases, these sRNAs are homologous to the target loci and not to either upstream or downstream regions. Genome-wide, *pyrF* and *pyrG* were among the genes most strongly differentially enriched for sRNAs in the epimutant versus the revertant strains, even without complete reversion to wild-type levels in the revertants (S1 Fig). Expression of the *pyrF* or *pyrG* specific sRNAs was lost upon reversion to 5-FOA sensitivity, although the E1 revertant did not return completely to parental levels after five passages.

**Fig 2.**
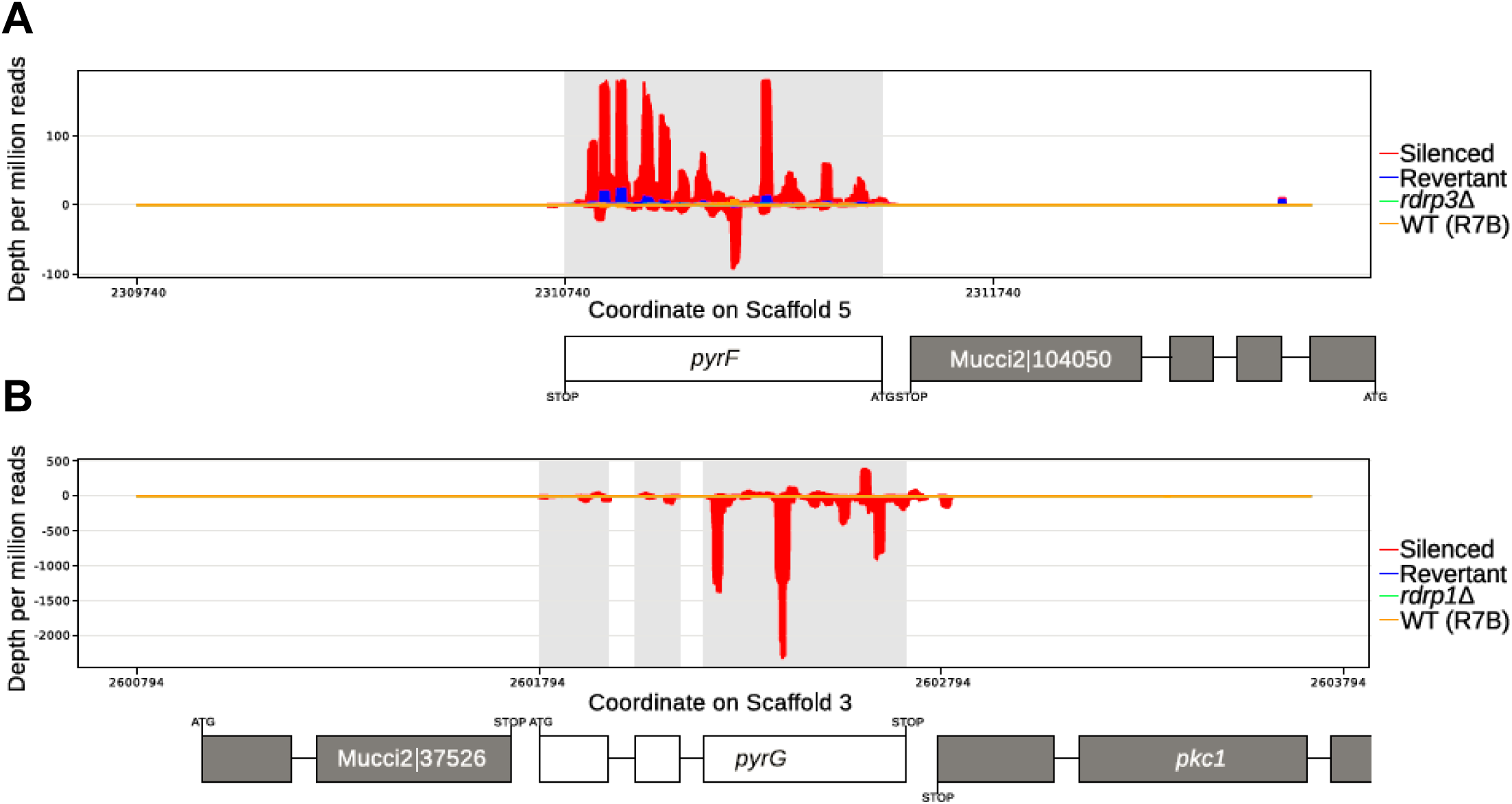
5-FOA resistance is associated with increased levels of sRNAs against either the *pyrF* or *pyrG* locus. **(A)** Representative diagram of sRNAs mapped across the *pyrF* locus showing accumulation of both sense (- values) and antisense sRNA (+ values) in epimutant E1 (in red). Expression levels are greatly decreased in the revertant after 5 passages without selection (in blue). No increase in sRNA levels is seen in the surrounding regions. **(B)** Representative diagram of sRNAs mapped across the *pyrG* locus showing accumulation of both sense (+ values) and antisense sRNA (- values) in epimutant E4 (in red), with greatly decreased sRNA levels in the revertant after 5 passages without selection (in blue). No increase of sRNA levels is observed in the surrounding regions.

### *pyrF* and *pyrG* epimutants harbor sRNAs with characteristic features

sRNAs from 5-FOA-resistant *pyrF* or *pyrG* epimutants also shared characteristics typical of sRNAs involved in the canonical RNAi pathway. These features included a high prevalence of a 5’ terminal uracil, which was found in antisense sRNAs in particular (Fig. 3a, 3b). Representative analysis from the *pyrF* epimutant E1 is shown here (Fig. 3a), as well as from the *pyrG* epimutant E4 (Fig. 3b). The same 5’ uracil predominance was observed in the few antisense sRNA reads found in the revertants; for better visualization a version of this figure with a scaled Y-axis has also been included (S2 Fig). This 5’ uracil prevalence was not identified in sense sRNAs from the same regions. In addition, the lengths of sRNA molecules homologous to these loci were predominantly between 21 and 24 nucleotides (Fig. 3c, 3d), a second feature of sRNAs generated by the canonical RNAi pathway and which interact with the RNAi effector protein Argonaute.

**Fig 3.**
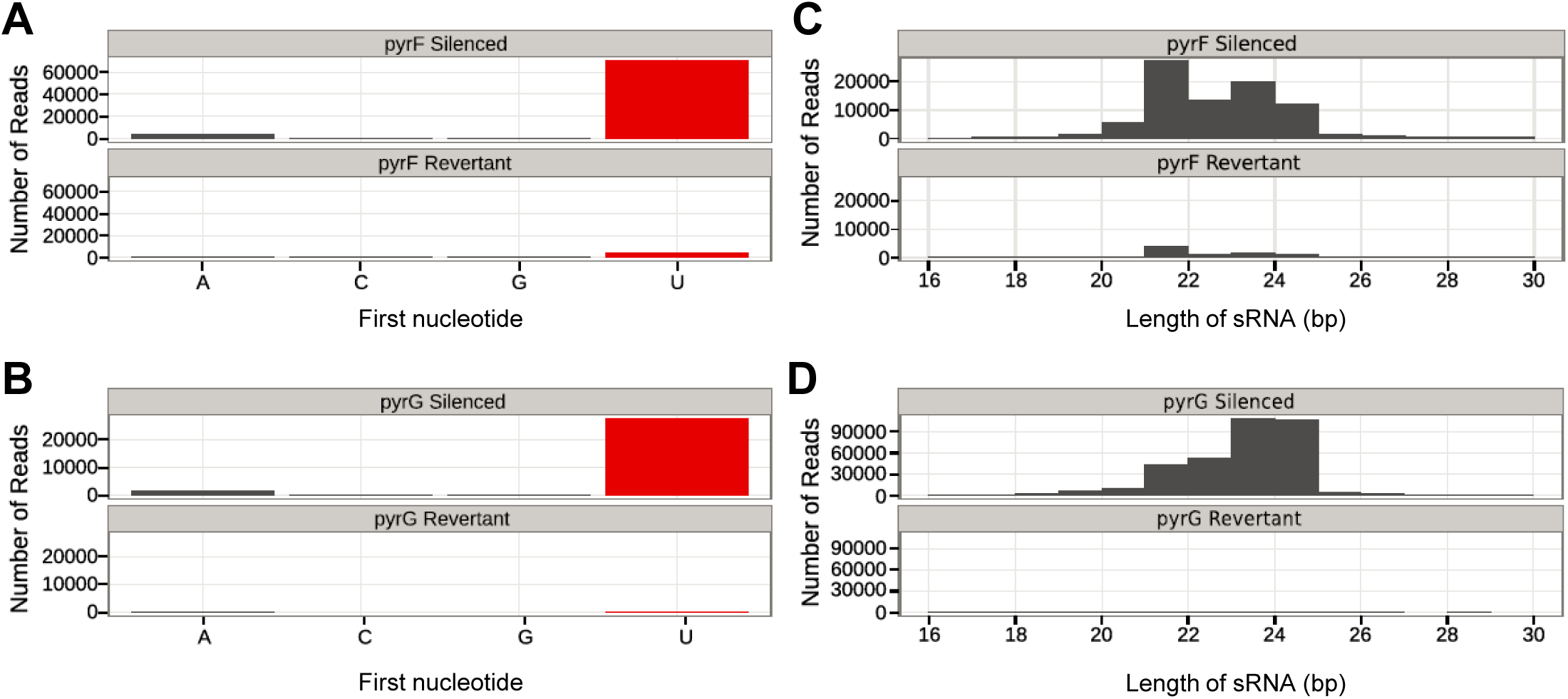
Length and terminal nucleotide analysis of sRNAs in epimutants. Antisense sRNAs from epimutant strains and their corresponding revertants that map against the *pyrF* and *pyrG* loci. **(A)** Analysis of the 5’ nucleotide of antisense sRNAs that map to the *pyrF* locus, isolated from the *pyrF* epimutant strain E1 and revertant. **(B)** Analysis of the 5’ nucleotide of antisense sRNAs that map to the *pyrG* locus, isolated from the *pyrG* epimutant strain E4 and revertant. **(C)** Analysis of the size of antisense sRNAs that map to *pyrF* in strain E1 and revertant. **(D)** Analysis of the size of antisense sRNAs that map to *pyrG* in strain E4 and revertant.

### A subset of genes with similarity to transposons exhibits altered sRNA levels

Analysis of genome-wide sRNA content also revealed a subset of genes that behaved unexpectedly in different samples. This set of genes had reduced sRNA content in the E2 epimutant, an *rdrp3*Δ mutant, compared to the rest of the *rdrp3*Δ strains that were sequenced (S3 Fig). Interestingly, while sRNA levels of these genes in the wild-type parent of the *rdrp1*Δ mutant were similar to levels in the *rdrp3*Δ mutant and its wild-type parent, all three sequences of *rdrp1*Δ strains in this study had lower sRNA levels corresponding to the same set of genes that behaved unusually in the E2 revertant (S3 Fig). A cutoff of 15-fold enrichment in the E4 revertant over the E4 epimutant was established, which selected 516 genes. Analysis of this gene set was complicated by generally low quality functional annotation of the *Mucor* genome. These genes were not grouped in any genomic location region but were relatively evenly distributed, appearing on every scaffold of the genome over 41 kb in size (S3 Fig). A search for conserved domains in this gene set revealed only 152 genes that encoded identifiable functional domains. However, 91 of these genes had predicted functions consistent with transposons or retrotransposons, including reverse transcriptase or transposase domains. These results may suggest that RdRP1 plays a role in repressing transposable elements via sRNA. However, the aberrant behavior of the E2 epimutant is not explained by this hypothesis because both the epimutant and its revertant are in the *rdrp3*Δ background. This suggests another level of regulation of this unusual class of sRNA.

### Epimutants exhibit reduced expression of target genes

Analysis of *pyrF* and *pyrG* mRNA expression levels by quantitative real-time PCR (qRT-PCR) showed a decrease in expression levels in epimutant isolates corresponding with sRNA generation. In *pyrF* epimutant strains, expression of *pyrF* mRNA was significantly decreased relative to expression in the *rdrp3* mutant parental background. Moreover, *pyrF* expression levels were restored upon reversion of the *pyrF* epimutation after five or ten passages (Fig. 4 a). As expected, no significant decrease was observed in *pyrG* expression in these *pyrF* epimutants either before or after reversion (Fig. 4b). Correspondingly, in the *pyrG* epimutant strain, decreased expression of *pyrG* but not *pyrF* mRNA was observed, with a subsequent increase upon reversion to 5-FOA sensitivity (Fig. 4 c, d).

**Fig 4.**
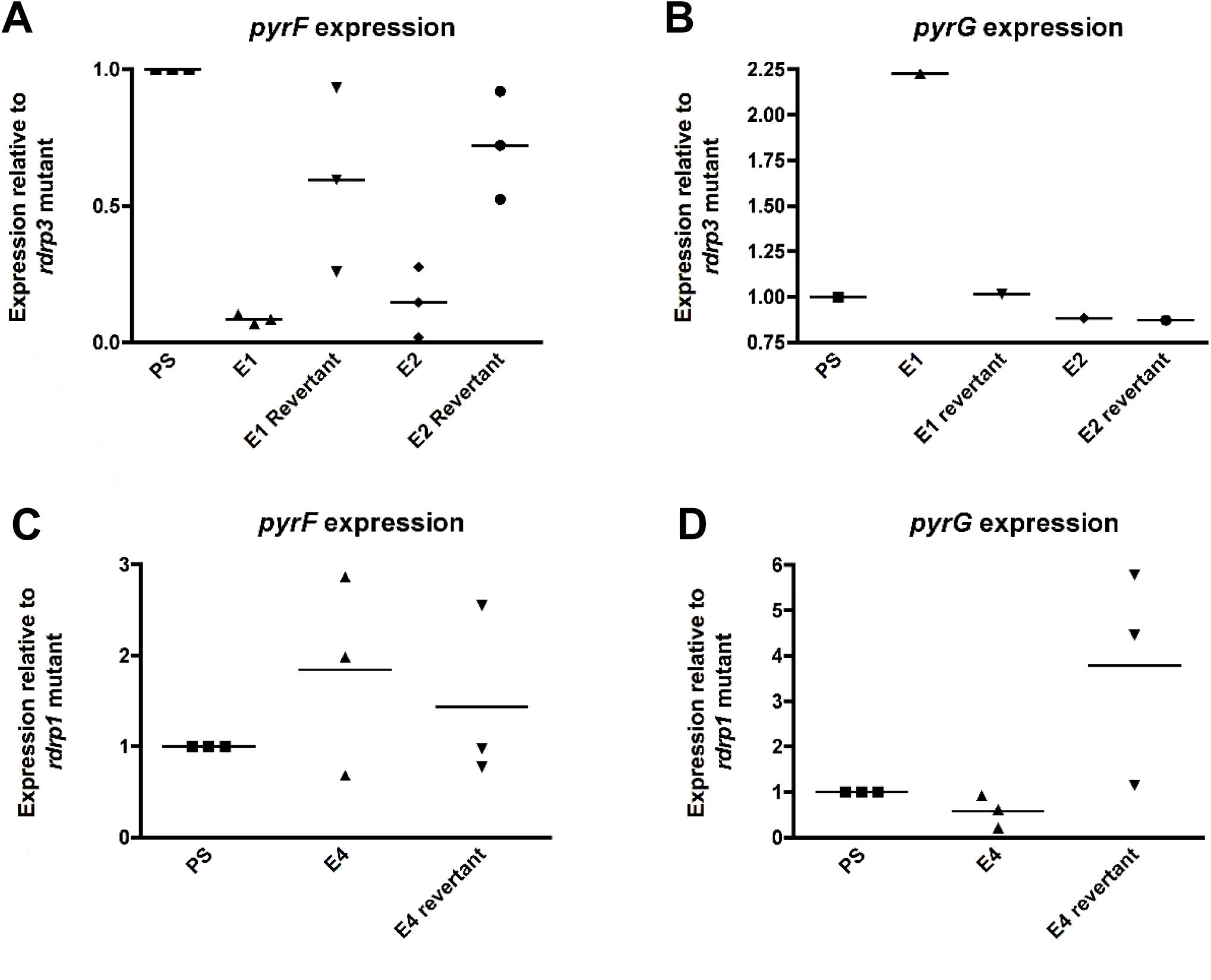
Epimutation decreases expression of *pyrF* and *pyrG* mRNA. **(A)** Expression of *pyrF* mRNA in *pyrF* epimutant and revertant strains (Passage 5, P5) as determined through qRT-PCR, with actin expression used for the reference gene. Gene expression levels were normalized relative to the *rdrp3*Δ parental strain (PS), using actin as the reference gene via the comparative ΔΔCt method. N=3 experimental replicates; error bars depict standard error of the mean (SEM). Significance determined via one-way ANOVA (P=0.0005, F=13.37, 4 degrees of freedom) with post-hoc Tukey’s Multiple Comparison test. **(B)** Expression of *pyrG* mRNA in *pyrF* epimutant and revertant strains. N=1. **(C)** Expression of *pyrF* mRNA in a *pyrG* epimutant and revertant strain as determined through qRT-PCR, with actin expression used for the reference gene. Percent expression was normalized relative to *rdrp1*Δ parental strain (PS). N = 3 experimental replicates; error bars depict SEM. Significance determined via one-way ANOVA (P=0.51, F=0.74, 2 degrees of freedom). **(D)** Expression of *pyrG* mRNA in *pyrG* epimutant and revertant strain. N = 3 experimental replicates; error bars depict SEM. Significance determined via one-way ANOVA (P=0.059, F=4.7, 2 degrees of freedom).

## Discussion

Epigenetic alteration of gene expression can lead to marked changes in phenotype across a variety of organisms. The phenomenon of epimutation was first described in plants and later in cancer biology; these particular alterations are attributable to extensive DNA methylation leading to gene silencing. Epimutations in snapdragons produce a phenotype wherein normal floral bilateral asymmetry is converted to radial symmetry [30]. In the field of cancer research, there is growing awareness that carcinogenesis can be driven by epimutation rather than mutations, including but not limited to cancers such as hereditary nonpolyposis colorectal cancer or BRCA-associated breast cancer [31-35]. Another role of epimutation that has gained attention is as a mechanism of drug resistance, with a particular focus on the roles played by DNA methylation and long noncoding RNAs in tumor drug resistance [36, 37]. Finally, a third form of epigenetic drug resistance, RNAi-dependent epimutation, was discovered to be a novel and transient mechanism of resistance to the agent FK506 in pathogenic fungi [13].

Identification of 5-FOA-resistant *Mucor* epimutants confirms that this mechanism is broader than had been previously demonstrated. Epimutation is capable of conferring resistance to multiple antifungal agents with different mechanisms of action, by targeting multiple genes. 5-FOA-resistant epimutant strains were identified that demonstrated silencing of either the *pyrF* locus or the *pyrG* locus. Therefore, generalization of the mechanism suggests that epimutation may broadly contribute to resistance by silencing a variety of drug target genes. No specific triggers for RNAi-based epimutation have been identified to date, although various stress conditions were previously tested [13]. The previous locus of epimutation, *fkbA*, was noted to have an overlapping gene (*patA*), but deletion of *patA* did not cause a loss of epimutation [13]. *pyrF* and *pyrG* do not have any overlapping flanking genes.

Rapid loss of silencing was observed in 5-FOA-resistant epimutant strains after five to ten passages without drug selection pressure. Epimutation – a transient and reversible phenomenon – may provide multiple advantages over genetic mutations that stably alter DNA sequence. In *Mucor*, which is aseptate and multinucleate, RNA-based silencing may induce more rapid and complete loss of function of disadvantageous genes compared to a recessive nuclear mutation, which would be required to sweep the population to become homokaryotic. In addition, the reversible nature of epimutation allows for subsequent reversal of adaptations that may be disadvantageous after a selective pressure is no longer present. For example, the uracil auxotrophy induced secondary to 5-FOA resistance could affect growth in low uracil conditions; under such conditions, epimutants, which can rapidly revert to wild-type and resume uracil synthesis, would have an advantage over *pyrF* or *pyrG* mutants. In support of this hypothesis we observed that *pyrF* and *pyrG* epimutants grew more effectively than a *pyrG* mutant strain in MMC lacking uracil supplementation, indicating that the epimutants may have incompletely silenced the *pyrF* or *pyrG* gene or may be undergoing reversion in response to selective pressure (Fig. 1). This is further supported by our qPCR data that demonstrated reduced, but not abolished, levels of *pyrF* or *pyrG* expression in the respective epimutants (Fig. 4).

The phenomenon of epimutation could thus be comparable to other described instances of fungal epigenetic heterogeneity, serving as a bet-hedging strategy that enables rapid and reversible responses to a variety of environmental conditions. One previously described example is telomeric silencing: genetic markers located near the telomeres of *S*. *cerevisiae*, including *URA3* as well as *ADE2*, were demonstrated to be variably silenced in a given population of yeast [38, 39]. These mixed populations (*URA*^+^*/ura*^-^or *ADE*^*+*^*/ade*^*-*^) are attributable to telomeric heterochromatin expanding and contracting across the integrated gene, resulting in silencing or expression. Likewise, in the fungal pathogen *C*. *neoformans*, the phenomena of sex-induced silencing or mitotic-induced silencing can be observed after tandem insertions of transgenes such as *ADE2*. The variable silencing of this tandem array can be observed through the phenomenon of variegation of colonies with both *ADE* and *ade-* phenotypes [19, 21, 40].

*Mucor* is known to possess multiple functional RNAi pathways [9, 41]. The sRNAs generated from the *pyrF* and *pyrG* loci show hallmark properties of RNAs that induce silencing through the core RNAi pathway [41]. Furthermore, sRNAs from *pyrG* localized to the exons of these genes, suggesting that the sRNAs were most likely generated and processed from mature mRNA. The gene *pyrF* contains no introns, but sRNAs were found to localize across the entire open reading frame without extending into neighboring regions. This suggests introns are not required for epimutation, and thus epimutation is distinct from previously described mechanisms of RNAi-mediated degradation that target poorly spliced introns [42].

5-FOA-resistant epimutants were discovered in two distinct genetic backgrounds: the *rdrp1* and *rdrp3* mutants, each of which lacks one of the three RNA-dependent RNA polymerases with roles in RNAi in *Mucor*. However, unlike the previous report of FK506-resistant epimutants, no 5-FOA-resistant epimutant strains were identified in wild-type strains or in an *r3b2Δ* strain mutated for a different RNAi component (S2 Table). In both the *rdrp1* and *rdrp3* mutant backgrounds, the overall frequency of 5-FOA-resistant epimutants was notably lower than the frequency of FK506-resistant epimutants, potentially due to the auxotrophic effect caused by loss of *pyrF* or *pyrG*. These RNAi deficient strains were previously demonstrated to have an increased frequency of epimutation relative to wild-type [13, 14]. Hence, one possibility is that the frequency of wild-type epimutants resistant to 5-FOA may be even lower that the frequency seen in RNAi mutant strains, making these wild-type epimutants difficult to isolate. Alternatively, it is possible that these mutant backgrounds are required for the isolation of 5-FOA-resistant epimutants. If RNAi deficiency is required for generation of 5-FOA-resistant epimutants, these findings would illustrate an interesting potential pathway to drug resistance that combines both a Mendelian (*rdrp1Δ* or *rdrp3*Δ) and an epigenetic factor in a two-step process.

Epimutation may enable an organism to temporarily resist environmental stresses to provide time for more permanent genetic diversity to arise. For example, induction of drug tolerance has been shown to play a role in subsequent mutation and the eventual development of bona fide drug resistance in bacteria [43, 44]. Similarly, aneuploidy has been reported to serve as a transient evolutionary adaptation that enables other genetic changes to arise [45]. One 5-FOA-resistant strain generated in this study, epimutant E7, initially expressed sRNA against *pyrF* and had no mutations in *pyrF* or *pyrG*. It lost this sRNA expression by passage 15, but did not revert to 5-FOA sensitivity even after 70 passages without selection (S4 Fig). Neither *pyrF* nor *pyrG* mutations were identified in this strain after passaging. One potential explanation based on these results is that epimutation provided transient relief from drug toxicity for this isolate and thus enabled the development of a more permanent form of resistance that remains to be elucidated.

In broader clinical terms, it is interesting to consider the role epimutation may play in *Mucor’s* intrinsic resistance to many antifungal agents, and whether epimutation may affect development of further resistance. For example, it has been suggested that amino acid substitutions in the *Mucor* Erg11/CYP51 enzyme, the target of the azole drug class, may explain part of *Mucor*’s innate resistance to certain structural classes of azoles (i.e. short-vs long-tailed azoles) [46]. However, this distinction alone is not sufficient to explain why only two azoles possess efficacy against *Mucor*, and additional mechanisms for intrinsic azole resistance should be investigated. In addition, epimutation could play a role in development of resistance to effective antifungals. The two front-line antifungals in clinical use against mucormycosis are the azole isavuconazole and the polyene amphotericin B. Resistance to azoles and polyenes in other pathogenic fungi, such as *Candida* species, can be mediated by loss of the ergosterol biosynthetic enzymes Erg3 and Erg6 [47-52]. Using bioinformatic analysis we have identified three candidate *ERG6* homologs and one candidate *ERG3* homolog in *Mucor* and we hypothesize that epimutation could induce silencing of these genes under appropriate drug selection, leading to acquired drug resistance. In particular, the presence of multiple copies of the gene encoding Erg6 could make this enzyme an especially appealing target for epimutation; if there is sufficient homology between these copies, we hypothesize RNAi could induce silencing of all three copies at once, instead of requiring mutations at all three loci to develop resistance.

Identification and characterization of 5-FOA resistance via RNAi-based epimutation advances understanding of the general mechanisms of drug resistance in *Mucor circinelloides*. The transient nature of epimutation is advantageous as it allows for rapid, facile reversion and flexible responses to changing conditions, such as uracil auxotrophy versus drug stress, enabling better adaptation to stressful conditions. Further questions that remain include whether RNAi-based epimutation occurs in other fungal species or other organisms with active RNAi systems. Further elucidation of the mechanism of epimutation advances our understanding of RNAi, drug resistance, and stress response mechanisms and may offer novel approaches to combat antifungal drug resistance.

## Materials and Methods

### Strains and growth conditions

All epimutants in this study were generated from strains of *Mucor circinelloides* forma *lusitanicus*. *M*. *circinelloides* f. *lusitanicus* RNAi mutant strains MU439, MU440, and MU500 (independently derived strains with the genotype *leuA-pyrG-rdrp3Δ*∷*pyrG*) and MU419 (*leuA-pyrG-rdrp1Δ*∷*pyrG*) were previously generated from the uracil and leucine auxotrophic strain MU402, which was in turn derived from the wild-type strain CBS277.49 [13, 14, 26]. As these four RNAi mutant strains were generated by using a functional copy of the *pyrG* gene to interrupt the target RNAi gene, each strain contains a mutant, nonfunctional copy of *pyrG* at the original locus as well as a functional copy inserted in an RNAi component gene. MU439, MU419, and the wild-type strain R7B served as controls for *M*. *circinelloides* f. *lusitanicus* studies, as appropriate. The strain 1006PhL was used for all *M*. *circinelloides* f. *circinelloides* studies.

Strains were grown at room temperature (approximately 24°C) with light exposure. Strains were cultured on MMC media at pH=4.5 (10 g/L casamino acids, 20 g/L glucose, and 0.5 g/L yeast nitrogen base without amino acids or ammonium sulfate) [26]. Media was supplemented with both uridine (0.061 g/L) and uracil (0.056 g/L) for potentially auxotrophic strains. 5-FOA selection was performed on MMC plates supplemented with uracil/uridine and 2.5 mg/mL 5-FOA. Passages were performed in liquid YPD (10 g/L yeast extract, 20 g/L peptone, 20 g/L dextrose) and on YPD agar.

### Generation and phenotypic analysis of 5-FOA-resistant mutants and epimutants

Epimutant candidates were generated by spotting *Mucor* spores on MMC media supplemented with 5-FOA and uridine/uracil; plates were incubated for approximately two weeks or until patches of resistant hyphal growth emerged from the periphery of drug-sensitive colonies, which were identified as colonies with severely stunted hyphae. Resistant isolates were passaged for at least two rounds of vegetative growth and sporulation under 5-FOA selection prior to sRNA analysis, to ensure a high proportion of drug resistance in the mycelia.

Epimutant strains were passaged in liquid YPD media without drug selection to induce reversion. For the first passage, spores were added to 3 mL of media and grown overnight at 30°C with shaking at 250 rpm. Subsequent passages were performed using a sterile wooden stick to break off a small portion of mycelia for transfer to fresh media. The final passage was performed using a sterile wooden stick to break off a small portion of mycelia that was placed on a YPD plate without drug selection; the plate was then incubated at room temperature (∼24°C) with light to allow for growth and sporulation. Spores were collected in sterile water for subsequent analyses.

### Nucleic acid extractions

Isolates were grown on MMC media, pH=4.5, supplemented with 2.5 mg/mL of 5-FOA and uridine/uracil as needed. DNA was extracted from hyphae using the MasterPure Yeast DNA Purification Kit (Epicenter Biotechnologies, Madison, WI), with the preliminary step of adding ∼100 µL of 425-600 µm glass beads and vortexing for one minute to break up hyphae. *pyrF* and *pyrG* were sequenced in all resistant isolates to rule out genetic mutations; primers are listed in S1 Table.

Isolates for RNA extraction were grown on plates overlaid with sterile cellulose film (ultraviolet irradiated for 10 minutes per side) to allow for easier removal of hyphae without agar contamination. Small and total RNAs were extracted using the mirVana kit (Ambion, Foster City, CA) for hybridization and qPCR analysis.

### sRNA hybridization

For sRNA hybridization, sRNA for each sample (3.5 µg) was separated by electrophoresis on 15% TRIS-urea gels, transferred to Hybond N+ filters, and cross-linked by ultraviolet irradiation as previously described (2 pulses at 1.2 x 10^5^µJ per cm^2^) [13]. Prehybridization was carried out using UltraHyb buffer (Ambion) at 65°C. *pyrF* and *pyrG* antisense-specific and 5s rRNA probes were prepared by *in vitro* transcription using the Maxiscript kit (Ambion); primers are listed in S1 Table. After synthesis, riboprobes were treated by alkaline hydrolysis as previously described [23], to generate an average final probe size of ∼50 nucleotides.

### mRNA quantification

Quantification of *pyrF* and *pyrG* mRNAs was performed by quantitative real-time PCR. Single-stranded cDNA was synthesized using AffinityScript (Stratagene, La Jolla, CA) from RNA samples treated with Turbo DNase (Ambion). cDNA synthesized without the RT/RNase enzyme mixture was used as a “no-RT control” to control for contamination by residual genomic DNA. Expression of target genes was measured using Brilliant III ultra-fast SYBR green QPCR mix (Stratagene) using an Applied Biosystems 7500 Real-time PCR system. Technical triplicates were performed for all samples in each run, and three biological replicates were performed for each experiment. Gene expression levels were normalized using actin as the reference gene via the comparative ΔΔCt method. Primers are listed in S1 Table.

## Statistics

One-way ANOVAs were used to determine the significance of qPCR replicates, with Tukey’s Multiple Comparison Test as a post-hoc test where appropriate. All statistical analysis was performed using GraphPad Prism.

### High-throughput sRNA sequencing and mapping

sRNA libraries were prepared and sequenced at the Duke Center for Genomic and Computational Biology using the Illumina TruSeq Small RNA Library Prep Kit coupled with agarose gel size selection for the miRNA library. Reads have been deposited at GEO under project accession number GSE113706.

Reads were trimmed using Trim Galore! with default settings to remove adapters [53]. Trimmed reads were then mapped to the *Mucor circinelloides* genome from the JGI using Bowtie [54, 55]. Reads mapping to gene loci were counted using Cufflinks and guided by genome annotation from the Joint Genome Institute (JGI) genome assembly [56].

## Acknowledgements

We thank Anna Floyd-Averette and Shelly Clancey for technical support, and Andrew Alspaugh, Silvia Calo, Shelly Clancey, Victoriano Garre, María Isabel Navarro-Mendoza, Francisco E. Nicolás, Carlos Pérez-Arques, Shelby Priest, Cecelia Wall, Santiago Torres Martinez, and Rosa Maria Ruiz Vazquez for critical reading of the manuscript.

## Supporting information

**S1 Fig. *pyrF* and *pyrG* are differentially enriched for sRNAs in epimutant vs. revertant strains.**

Genome-wide sRNA levels are plotted between two sequenced libraries, with values for one library plotted on the X and the other on the Y. The point representing *pyrF* is depicted in red and the *pyrG* is depicted in blue. E1 and E2 are *pyrF* epimutant strains; E4 is a *pyrG* epimutant strain.

**S2 Fig. Epimutant revertants demonstrate a 5’ uracil predominance in antisense sRNAs against *pyrF* and *pyrG*.**

Analysis of the 5’ nucleotide of antisense sRNAs that map to the *pyrF* and *pyrG* loci. Data from Fig. 3 is replotted here with an expanded y-axis to enable easier comparison of 5’ terminal nucleotides in sRNA from revertant strains. **(A)** 5’ terminal nucleotides of antisense sRNAs isolated from *pyrF* epimutant E1 and revertant. **(B)** 5’ terminal nucleotides of antisense sRNAs isolated from *pyrG* epimutant E4 and revertant.

**S3 Fig. A set of genes with discordant sRNA expression in epimutants vs. revertants are evenly distributed across the *Mucor* genome.**

**(A)** Genome-wide sRNA levels are shown with the gene set that is expressed more than 15-fold higher in the E2 revertant than in the E2 epimutant shaded in red. That same gene set is also shaded in the comparison of the *rdrp1*Δ parent strain with the WT parent to demonstrate that the same gene set is behaving anomalously in both comparisons. **(B)** Genes with discordant sRNA expression are shown across the *Mucor* genome (red bars not to scale relative to scaffold). These genes appear on every scaffold of the genome that is greater than 41 kb in size.

**S4 Fig. Epimutant E7 maintains 5-FOA resistance after cessation of sRNA expression.**

**(A)** The *pyrF* epimutant E7 maintains a degree of 5-FOA resistance even after 70 passages without drug selection (P70). An independent repeat of passaging demonstrates continued 5-FOA resistance through 40 passages (P40v2). Strains were grown on MMC media, MMC supplemented with uridine and uracil, and MMC supplemented with 5-FOA, uridine, and uracil. P, *rdrp1Δ* parental strain (MU419). **(B)** sRNA hybridization of passaged strains of epimutant E7. Epimutant E7 expresses sRNA against *pyrF*, but this sRNA is no longer expressed from 15 passages (P15) through 70 passages (P70). Similarly, strains from an independent set of passages demonstrate no sRNA against *pyrF* at passages 15 (P15v2), 30 (P30v2), or 40 (P40v2). The top portion of the gel was stained with ethidium bromide (EtBr) to visualize the 5S rRNA loading control, after which sRNA was transferred to a membrane for hybridization with an antisense-specific probe against *pyrF*.

**S1 Table. Primers used in this study**

**S2 Table. Strains generated in this study**

