## Supplementary material for "Broad antifungal resistance mediated by RNAi-dependent epimutation in the basal human fungal pathogen *Mucor circinelloides*"

### Supplemental Figure S1

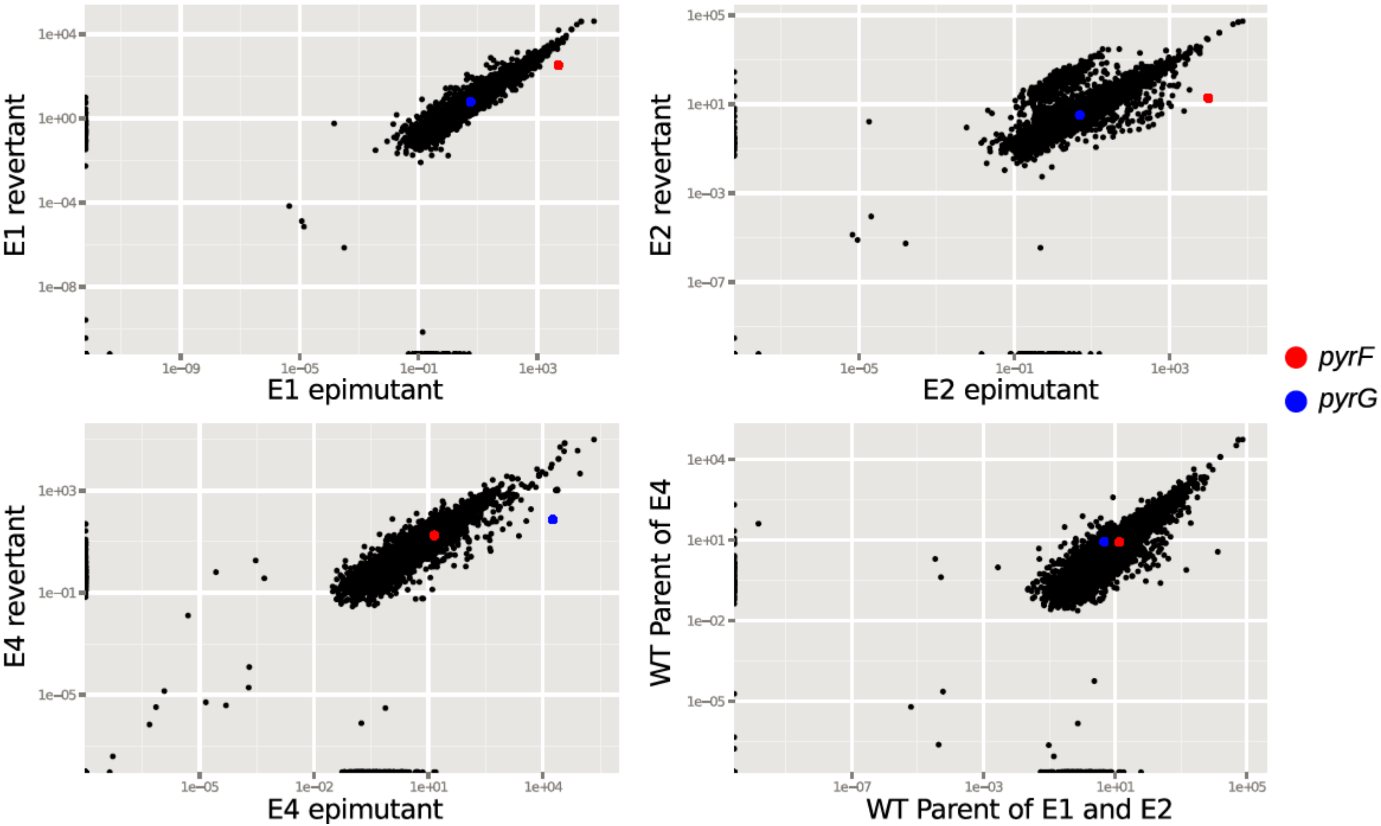

### Supplemental Figure S2

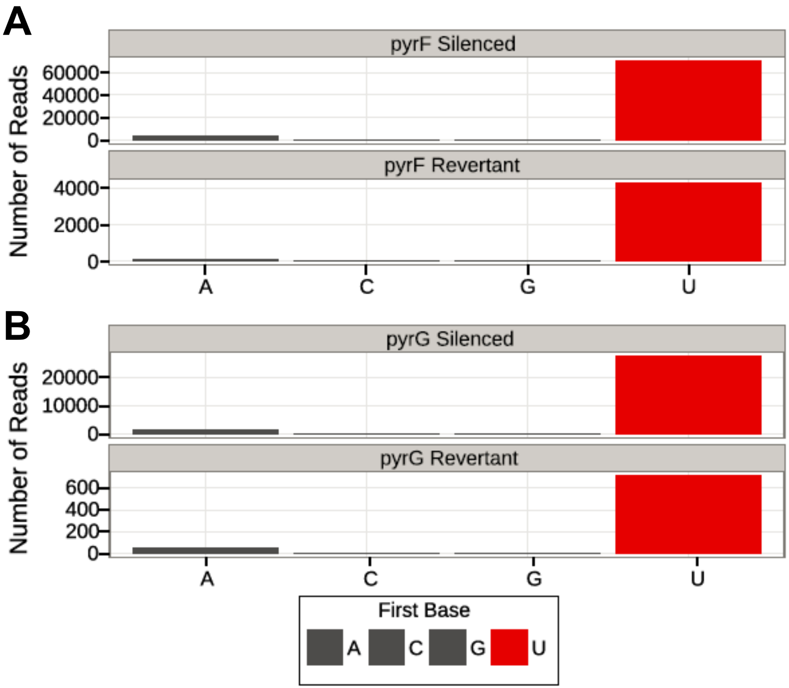

### Supplemental Figure S3

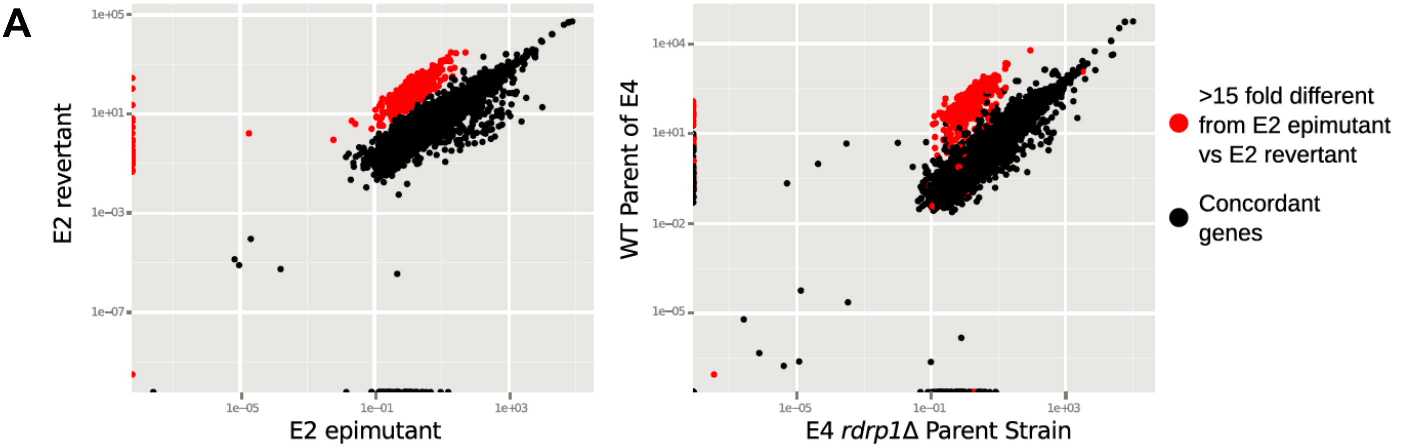


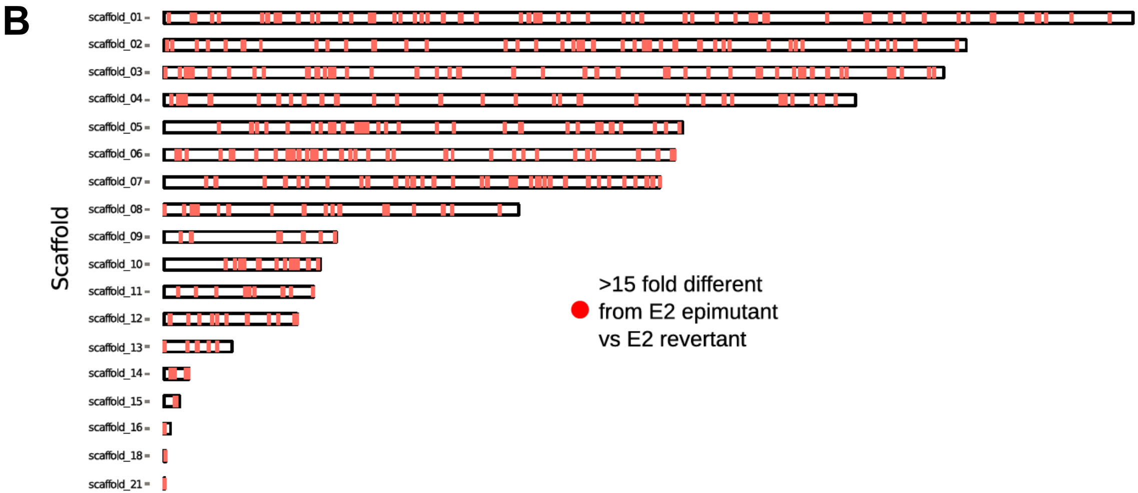

### Supplemental Figure S4

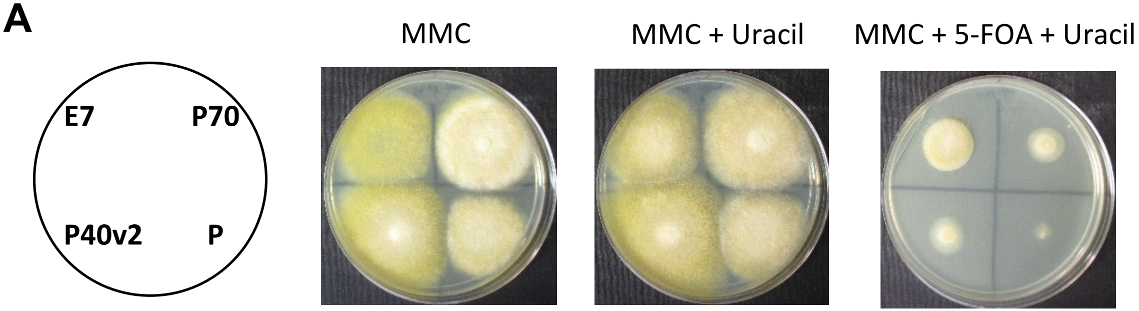


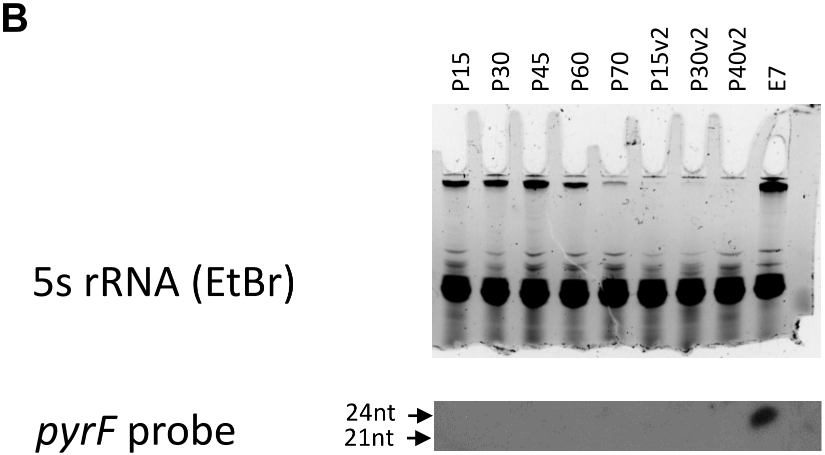

#### S1 Table. Primers used in this study

| Name | Sequence | Use |
|---|---|---|
| **JOHE41713** | GAGCTGCTGTAAGGCTGGAC | R7B *pyrF* forward |
| **JOHE41714** | TGAAATGGGATCCATCAACA | R7B *pyrF* reverse |
| **JOHE41962** | AGAGTCACCCATTTTGAGTGC | R7B *pyrF* forward |
| **JOHE41963** | TTGCTAGATCCGGGTTTCAC | R7B *pyrF* reverse |
| **JOHE42127** | CATTTGGCACCATCATTCAG | Locus-specific amplification of *pyrG* in *rdrp3* mutant strains |
| **JOHE42128** | GTTGGGTCGACTGTCGTTTT | Locus-specific amplification of *pyrG* in *rdrp3* mutant strains |
| **JOHE43751** | CGTCAGATCGTTCTTGCAGG | Locus-specific amplification of *pyrG* in *rdrp1* mutant strains |
| **JOHE43752** | CGGTCTCTCCCCATAACACG | Locus-specific amplification of *pyrG* in *rdrp1* mutant strains |
| **JOHE44080** | ATTGGGATGCTGTTGTCCAC | R7B *pyrG* sequencing |
| **JOHE42081** | CAAATGACATTAGCCCCTTGA | R7B *pyrG* sequencing |
| **JOHE42082** | CGAGGTTGGTCTTCCTCTTG | R7B *pyrG* sequencing |
| **JOHE42083** | TGGTCTGGGTTGCCATAGAT | R7B *pyrG* sequencing |
| **JOHE41665** | TTGAGTGTGCGGAGATCTTG | 1006PhL *pyrF* forward |
| **JOHE41666** | TGACCTCACGTGGTTGATCT | 1006PhL *pyrF* reverse |
| **JOHE41667** | CGCTGGATCTGGGTGATATT | 1006PhL *pyrG* forward |
| **JOHE41668** | CCTGCCTTAACTCCCATCAA | 1006PhL *pyrG* reverse |
| **JOHE41669** | CGAGGACTTTGACCGTGATT | 1006PhL *pyrG* sequencing |
| **JOHE38278** | GAGGAATGAGACCGGGGTAACCAC | 24mer for size standard on sRNA blots |
| **JOHE42163** | AGATCCACGATCACGAGATGA | 21mer for size standard on sRNA blots |
| **JOHE42212** | TAATACGACTCACTATAGGGTTGAGTG TGCGGAGATCTTG | T7 promoter and 1006PhL *pyrF* forward. For sRNA antisense probe synthesis combined with JOHE41666 |
| **JOHE42354** | TAATACGACTCACTATAGGGCGCTGGA TCTGGGTGATATT | T7 promoter and 1006PhL *pyrG* forward. For sRNA antisense probe synthesis combined with JOHE41668 |
| **JOHE42440** | TAATACGACTCACTATAGGGAGAGTCA CCCATTTTGAGTGC | T7 promoter and *pyrF* forward primer, R7B. For sRNA antisense probe synthesis combined with JOHE41963. |
| **JOHE42441** | TAATACGACTCACTATAGGGCGAATGG AAAGTGAGTGGGT | T7 promoter and *pyrG* forward primer, R7B. For sRNA antisense probe synthesis combined with JOHE20867. |
| **JOHE20867** | GTACACTGGCCATGCTATCG | *pyrG* reverse primer, R7B |
| **JOHE37682** | TAATACGACTCACTATAGGGAGCTACG GCCATACAATGTTG | T7 promoter and 5s rRNA, for amplification of the 5S rRNA probe |
| **JOHE37683** | TAATACGACTCACTATAGGGGAACTAC AGCAACCAGTATTCCCA | T7 promoter and 5s rRNA, for amplification of the 5S rRNA probe |
| **JOHE42636** | ACCGCAAGGAAAAGAAGGAT | R7B *pyrF* qRT-PCR |
| **JOHE42637** | CAAGGACACCAGCAAGTTGA | R7B *pyrF* qRT-PCR |
| **JOHE44065** | TTGATGGAGCGTAAGCAATC | R7B *pyrG* qRT-PCR |
| **JOHE44066** | AGCAACCAAATCGTGGTCA | R7B *pyrG* qRT-PCR |
| **JOHE24077** | AAGCCCAATCCAAGAGAGGT | R7B actin qRT-PCR |
| **JOHE24078** | GCCTCAGTCAAGAGGACAGG | R7B actin qRT-PCR |

#### S2 Table. Strains generated in this study

| Strain | # | Epimutant | Genetic background | Mutation/cause of resistance |
|---|---|---|---|---|
| ***rdrp3*Δ** | **1** | | MU500 | Unknown; no mutations or *pyrF/pyrG* sRNA found |
| | **2** | | MU500 | Unknown; no mutations or *pyrF/pyrG* sRNA found |
| | **3** | | MU500 | Unknown; no mutations or *pyrF/pyrG* sRNA found |
| | **4** | | MU500 | Unknown; no mutations or *pyrF/pyrG* sRNA found |
| | **5** | | MU500 | Unknown; no mutations or *pyrF/pyrG* sRNA found |
| | **6** | | MU500 | Unknown; no mutations or *pyrF/pyrG* sRNA found |
| | **7** | | MU500 | Unknown; no mutations or *pyrF/pyrG* sRNA found |
| | **8** | | MU500 | Unknown; no mutations or *pyrF/pyrG* sRNA found |
| | **9** | | MU440 | Unknown; no mutations or *pyrF/pyrG* sRNA found |
| | **10** | | MU440 | Unknown; no mutations or *pyrF/pyrG* sRNA found |
| | **11** | | MU440 | Unknown; no mutations or *pyrF/pyrG* sRNA found |
| | **12** | **E1** | MU439 | ***pyrF* epimutant** |
| | **13** | | MU439 | Unknown; no mutations or *pyrF/pyrG* sRNA found |
| | **17** | **E2** | MU439 | ***pyrF* epimutant** |
| ***rdrp1*Δ** | **1** | **E6** | MU419 | ***pyrF* epimutant** |
| | **2** | | MU419 | *pyrG* G413A; exon splice site |
| | **3** | | MU419 | *pyrG* G413A; exon splice site |
| | **4** | | MU419 | *pyrG* G413A; exon splice site |
| | **5** | | MU419 | *pyrG* G413A; exon splice site |
| | **6** | | MU419 | *pyrG* G413A; exon splice site |
| | **7** | | MU419 | *pyrG* G413A; exon splice site |
| | **8** | | MU419 | *pyrG* G413A; exon splice site |
| | **9** | **E7** | MU419 | ***pyrF* epimutant** |
| | **10** | | MU419 | *pyrG* G413A; exon splice site |
| | **11** | | MU419 | *pyrG* G413A; exon splice site |
| | **12** | **E3** | MU419 | ***pyrF* epimutant** |
| | **13** | | MU419 | Unknown; no mutations or sRNA found |
| | **14** | **E4** | MU419 | ***pyrG* epimutant** |
| | **15** | | MU419 | Unknown; no mutations or sRNA found |
| | **16** | | MU419 | *pyrG* G413A; exon splice site |
| | **17** | | MU419 | *pyrG* G413A; exon splice site |
| | **18** | | MU419 | *pyrG* G413A; exon splice site |
| | **19** | | MU419 | *pyrG* G413A; exon splice site |
| | **20** | | MU419 | *pyrG* G413A; exon splice site |
| | **21** | | MU419 | *pyrG* G413A; exon splice site |
| | **22** | | MU419 | *pyrG* G413A; exon splice site |
| | **23** | **E5** | MU419 | ***pyrF* epimutant** |
| | **24** | | MU419 | *pyrG* G413A; exon splice site |
| | **25** | | MU419 | *pyrG* G413A; exon splice site |
| | **26** | | MU419 | *pyrG* G413A; exon splice site |

| | | | | |
|---|---|---|---|---|
| | **27** | | MU419 | *pyrG* G413A; exon splice site |
| ***r3b2*∆** | **1** | | MU412 | *pyrG* G413A; exon splice site |
| | **2** | | MU412 | *pyrG* G413A; exon splice site |
| | **3** | | MU412 | *pyrG* G413A; exon splice site |
| | **4** | | MU412 | Unknown; no mutations or sRNA found |
| | **5** | | MU412 | *pyrG* G413A; exon splice site |
| | **6** | | MU412 | *pyrG* G413A; exon splice site |
| | **7** | | MU412 | *pyrG* G413A; exon splice site |
| | **8** | | MU412 | *pyrG* G413A; exon splice site |
| | **9** | | MU412 | *pyrG* G413A; exon splice site |
| | **10** | | MU412 | *pyrG* G413A; exon splice site |
| | **11** | | MU412 | *pyrG* G413A; exon splice site |
| | **12** | | MU412 | *pyrG* G413A; exon splice site |
| | **13** | | MU412 | *pyrG* G413A; exon splice site |
| | **14** | | MU412 | Unknown; no mutations or sRNA found |
| **R7B** | **1** | | R7B | Unknown; no mutations or *pyrF/pyrG* sRNA found |
| | **2** | | R7B | Unknown; no mutations or *pyrF/pyrG* sRNA found |
| **1006PhL** | **1** | | 1006PhL | Unknown; no mutations or *pyrF/pyrG* sRNA found |
| | **2** | | 1006PhL | Unknown; no mutations or *pyrF/pyrG* sRNA found |
| | **3** | | 1006PhL | Unknown; no mutations or *pyrF/pyrG* sRNA found |
| | **4** | | 1006PhL | Unknown; no mutations or *pyrF/pyrG* sRNA found |
| | **5** | | 1006PhL | Unknown; no mutations or *pyrF/pyrG* sRNA found |
| | **6** | | 1006PhL | Unknown; no mutations or *pyrF/pyrG* sRNA found |
| | **7** | | 1006PhL | Unknown; no mutations or *pyrF/pyrG* sRNA found |
| | **8** | | 1006PhL | Unknown; no mutations or *pyrF/pyrG* sRNA found |
| | **9** | | 1006PhL | Unknown; no mutations or *pyrF/pyrG* sRNA found |
| | **10** | | 1006PhL | Unknown; no mutations or *pyrF/pyrG* sRNA found |
| | **11** | | 1006PhL | *pyrF* C535T (nonsense mutation) * |
| | **12** | | 1006PhL | *pyrF* C535T (nonsense mutation) * |
| | **13** | | 1006PhL | *pyrF* C535T (nonsense mutation) * |
| | **14** | | 1006PhL | *pyrF* C535T (nonsense mutation) * |
| | **15** | | 1006PhL | *pyrF* C535T (nonsense mutation) * |
| | **16** | | 1006PhL | *pyrF* C535T (nonsense mutation) * |
| | **17** | | 1006PhL | *pyrF* C535T (nonsense mutation) * |
| | **18** | | 1006PhL | Unknown; no mutations or *pyrF/pyrG* sRNA found |
| | **19** | | 1006PhL | Unknown; no mutations or *pyrF/pyrG* sRNA found |
| | **20** | | 1006PhL | Unknown; no mutations or *pyrF/pyrG* sRNA found |
| | **21** | | 1006PhL | Unknown; no mutations or *pyrF/pyrG* sRNA found |
| | **22** | | 1006PhL | Unknown; no mutations or *pyrF/pyrG* sRNA found |
| | **23** | | 1006PhL | *pyrF* C535T (nonsense mutation) * |
| | **24** | | 1006PhL | *pyrF* C535T (nonsense mutation) * |
| | **25** | | 1006PhL | *pyrF* C535T (nonsense mutation) * |
| | **26** | | 1006PhL | *pyrF* C535T (nonsense mutation) * |

| | | | |
|---|---|---|---|
| **27** | | 1006PhL | Unknown; no mutations or *pyrF/pyrG* sRNA found |
| **28** | | 1006PhL | Unknown; no mutations or *pyrF/pyrG* sRNA found |
| **29** | | 1006PhL | Unknown; no mutations or *pyrF/pyrG* sRNA found |
| **30** | | 1006PhL | Unknown; no mutations or *pyrF/pyrG* sRNA found |
| **31** | | 1006PhL | Unknown; no mutations or *pyrF/pyrG* sRNA found |
| **32** | | 1006PhL | Unknown; no mutations or *pyrF/pyrG* sRNA found |
| **33** | | 1006PhL | Unknown; no mutations or *pyrF/pyrG* sRNA found |
| **34** | | 1006PhL | Unknown; no mutations or *pyrF/pyrG* sRNA found |
| **35** | | 1006PhL | Unknown; no mutations or *pyrF/pyrG* sRNA found |
| **36** | | 1006PhL | Unknown; no mutations or *pyrF/pyrG* sRNA found |
| **37** | | 1006PhL | Unknown; no mutations or *pyrF/pyrG* sRNA found |
| **38** | | 1006PhL | Unknown; no mutations or *pyrF/pyrG* sRNA found |
| **39** | | 1006PhL | Unknown; no mutations or *pyrF/pyrG* sRNA found |
| **40** | | 1006PhL | Unknown; no mutations or *pyrF/pyrG* sRNA found |
| **41** | | 1006PhL | Unknown; no mutations or *pyrF/pyrG* sRNA found |
| **42** | | 1006PhL | Unknown; no mutations or *pyrF/pyrG* sRNA found |
| **43** | | 1006PhL | Unknown; no mutations or *pyrF/pyrG* sRNA found |
| **44** | | 1006PhL | *pyrF* C535T (nonsense mutation) * |
| **45** | | 1006PhL | *pyrF* C535T (nonsense mutation) * |
| **46** | | 1006PhL | *pyrF* C535T (nonsense mutation) * |
| **47** | | 1006PhL | *pyrF* C535T (nonsense mutation) * |
| **48** | | 1006PhL | *pyrF* C535T (nonsense mutation) * |
| **49** | | 1006PhL | *pyrF* C535T (nonsense mutation) * |
| **50** | | 1006PhL | *pyrF* C535T (nonsense mutation) * |
| **51** | | 1006PhL | *pyrF* C535T (nonsense mutation) * |
| **52** | | 1006PhL | *pyrF* C535T (nonsense mutation) * |
| **53** | | 1006PhL | *pyrF* T288 1bp deletion |
| **54** | | 1006PhL | *pyrF* C535T (nonsense mutation) * |
| **55** | | 1006PhL | Unknown; no mutations or *pyrF/pyrG* sRNA found |
| **56** | | 1006PhL | *pyrF* C535T (nonsense mutation) * |
| **57** | | 1006PhL | *pyrF* C535T (nonsense mutation) * |
| **58** | | 1006PhL | *pyrF* C535T (nonsense mutation) * |
| **59** | | 1006PhL | *pyrF* C535T (nonsense mutation) * |
| **60** | | 1006PhL | *pyrF* C535T (nonsense mutation) * |
| **61** | | 1006PhL | *pyrF* C535T (nonsense mutation) * |

* The prevalence of this single mutation in the 1006PhL background suggests these could potentially be sibling strains derived from a mutation present in the original population.
